# Characterizing Transition State in Mouse Vigilance with EEG–EMG Hypnodensity

**DOI:** 10.64898/2026.07.30.741729

**Authors:** Sadegh Rahimi, Monika Vadkertiova, Leesa Joyce, Andre Sevenius Nilsen, Carlo Mejia, Svenja L Kreis, Andreas Lieb, Taro Tezuka, Matteo Cesari, Thomas Fenzl

**Affiliations:** Institute for Pharmacology, Medical University of Innsbruck, 6020-Innsbruck, Austria; Department of Anesthesiology and Intensive Care, Technical University of Munich School of Medicine and Health, Munich, Germany; Institute of Basic Medical Sciences, University of Oslo, Norway; Faculty of Engineering, Information and Systems, University of Tsukuba, Japan; Department of Neurology, Medical University of Innsbruck, Innsbruck, Austria

**Keywords:** Hypnodensity, transition, intermediate state, EEG, machine learning

## Abstract

**Study Objectives:** Vigilance-state transitions are continuous biological processes, yet conventional rodent sleep scoring relies on discrete epochs that obscure intermediate states. As no standardized framework exists for characterizing these intermediate states in rodents, this study aimed to characterize the temporal dynamics of transitions in mice and validate a machine-learning approach for objective detection.

**Methods:** Chronic EEG and EMG recordings were obtained from male C57BL/6N mice. We extracted 56-second windows containing stable transitions between Wakefulness (WAKE), Non-Rapid Eye Movement Sleep (NREMS), and Rapid Eye Movement Sleep (REMS). Eight trained experts manually annotated the onset and duration of transitions to establish ground truth and assess inter-rater reliability. Using quantitative EEG/EMG features (e.g., spectral power, complexity, EMG variance) derived from stable states, Support Vector Machine (SVM) classifiers were trained to predict transition midpoints in independent test animals.

**Results:** Inter-rater agreement among experts was moderate to low, particularly for WAKE to NREMS and NREMS to REMS transitions, reflecting inherent ambiguity in manual scoring. Temporal analysis revealed distinct dynamics across transition types; NREMS to REMS transitions were significantly longer than all others, while REMS to NREMS transitions were the most abrupt. Despite the variability in human scoring, SVM models trained only on stable-state features successfully predicted expert-defined transition midpoints.

**Conclusions:** Our approach not only characterized the recognizable dynamics across transition types in mice, but also provides a reproducible framework for quantifying sleep-wake transitions, which is crucial for studying arousal stability and related impairments in disease.

**Statement of Significance:** Traditional sleep scoring enforces discrete boundaries between vigilance states, overlooking transitional dynamics that may be critical for understanding arousal regulation. We developed a novel hypnodensity-based framework to systematically identify and characterize intermediate vigilance states in mice using EEG-EMG recordings. By combining expert annotations with machine learning, we revealed that transitions between sleep and wake involve continuous processes with mixed state features, rather than instantaneous switches. This approach provides the first standardized method for quantifying transitional vigilance states in rodents, enabling deeper investigation of arousal instability in neurological disorders. Our framework advances automated sleep analysis beyond classical three-state classification

## Introduction

Sleep–wake regulation in mammals is a dynamic process governed by complex neural networks that alternate between vigilance states, including wakefulness (WAKE), non-rapid eye movement sleep (NREMS) and rapid eye movement sleep (REMS). These states are typically identified through electroencephalogram (EEG) and electromyogram (EMG) recordings. During wakefulness, the EEG typically exhibits low-amplitude, high-frequency activity accompanied by high EMG tone ^1^. In NREMS, the EEG is dominated by high-amplitude, low-frequency oscillations, with a reduced and variable EMG tone^2^. REMS, in contrast, shows a regular, low-amplitude, high-frequency EEG pattern combined with an almost complete absence of EMG tone^3^. In addition to three classical vigilance states, some studies in rodents have identified finer subdivisions of vigilance states, including distinctions between active and quiet wakefulness^4^, substates within NREMS^5^, and a pre-REM state^6^.

The well-known “flip-flop” model describes how mutually inhibitory sleep- and wake-promoting neuronal populations ensure rapid and stable transitions between vigilance states, preventing intermediate or unstable phases^7^. However, vigilance states are not strictly discrete and transitions between sleep and wake might be continuous processes that can involve partial activation or deactivation of specific neuronal networks^8,9^, rather than instantaneous switches. During these transitions, mixed features of sleep and wake may coexist, producing epochs that are difficult to classify with conventional frameworks^6,9^. Yet because classical scoring enforces artificial boundaries between states, these transitional dynamics remain largely unexplored.

From a dynamical systems perspective, these transitions represent trajectories between distinct attractor states^10,11^. Pinpointing the precise ’point of no return’ where neural activity shifts toward a new state is critical for mapping the underlying circuit bifurcations. While traditional epoch-based scoring provides a coarse overview of stability, it lacks the temporal resolution required to align transition dynamics with the rapid firing patterns of state-specific neuronal populations. The topic of sleep states and transitions between vigilance states may thus be a valuable area of research. But to achieve this, among others, better labeling and identification of changes in the state space is necessary.

In addition, understanding these intermediate vigilance states is crucial, as instability or abnormal timing of transitions has been implicated in several neuropsychiatric and neurological conditions. For instance, altered transition dynamics have been linked to abnormal arousal regulation in narcolepsy^12^ and increased seizure susceptibility in epilepsy^5^. Thus, quantifying how and when the brain switches between vigilance states may provide new insights into the mechanisms governing arousal stability and its impairment in disease.

While the American Academy of Sleep Medicine provides detailed guidelines for managing ambiguous or mixed epochs in human sleep scoring^13^, no equivalent standard exists for rodent data. Moreover, recent advances in automated sleep classification using machine learning have primarily focused on stable vigilance states (WAKE, NREMS, REMS) and largely overlooked the transitions that connect them^14^. Consequently, there is still no established framework to identify intermediate vigilance states in animal models for future usage in advanced sleep/wake analyses.

To address this gap, we sought to characterize intermediate vigilance states in mice using a combination of expert annotations and machine-learning methods. By asking 8 trained experts to mark transition periods and comparing their annotations, we quantified inter-rater reliability and identified the temporal structure of transitions. To capture the inherent ambiguity in sleep stage classification—where epochs often exhibit mixed characteristics—we adopted the hypnodensity approach, commonly used in human sleep research. This method represents each epoch as a probability distribution across sleep stages rather than forcing a single discrete label, thereby reflecting inter-scorer variability and epoch-level uncertainty^15,16^. Based on these expert-derived labels, we then trained support vector machine (SVM) classifiers using EEG and EMG features to predict the transition midpoints. We hypothesized that such models could reproduce expert-level accuracy, thereby providing an objective, reproducible framework for sleep–wake transitions in rodents.

## Methods

### Animals

Eleven male C57BL/6N mice (Charles River Laboratories GmbH, Germany), 12–18 weeks old (body weight 24–27 g), were used in this study. Animals were housed individually under a 12:12 h light– dark cycle (lights on/off: 09:00/21:00; temperature 22 ± 2 °C; humidity 55 ± 10%) with *ad libitum* access to food and water. All mice were maintained under identical environmental conditions. To reduce phase variability and ensure circadian alignment, mice were transferred from the animal facility to the laboratory seven days before surgery during the maintenance period (08:00 am to 09:00 am, lights ON). From one day before surgery until 4 days after surgery, mice received analgesia (carprofen) via drinking water (10 mg/kg bodyweight). All procedures were approved by the Committee of Animal Health Care of the State of Upper Bavaria, Germany (ROB-55.2–2532.Vet_02– 19–121) and conducted in accordance with EU guidelines for laboratory animal care and the ARRIVE guidelines^17^. The study was not preregistered on the Open Science Framework.

### Surgery and electrode implantation

For the electrode implantation, anesthesia induction was performed in an acrylic glass chamber with 4 Vol.-% isoflurane (CP-Pharma Handelsgesellschaft GmbH, Germany). Throughout the surgery, anesthesia was maintained at 1.8–2.0 Vol.-% isoflurane with a flow rate of 192 ml/min. A body temperature of 37°C was maintained using a homeothermic monitoring system (Harvard Apparatus, USA). For analgesia, Carprofen (4 mg/kg BW, Zoetis, Germany) was subcutaneously injected before surgery, and Lidocaine Hydrochloride (2%, bela-pharm GmbH & Co. KG, Germany) was applied to the incisions during surgery. After shaving the head, skin incisions were made to expose the upper cranium, and the periosteum was removed at surgery sites. One epidural EEG electrode (left occipital cortex, AP,Q−Q2.80 mm, ML,Q−Q2.41mm), reference electrode (AP, +0.62 mm, ML, −3.02 mm), and one EMG electrode (nuchal muscle) were implanted chronically. A printed circuit board socket (Preci-Dip, series 861, Delémont, Switzerland) holding all the electrodes was mounted and fixed to the cranium using dental cement (Paladur, Kulzer GmbH, Germany) and two jeweller’s screws (Ø 1.2Q×Q2 mm; Paul Korth GmbH, Lüdenscheid, Germany). The EEG and EMG electrodes were made of gold wires ( 150 µm, Haefner & Krullmann GmbH, Germany).

### Data acquisition

After surgery, the mice were allowed to recover for 10 days. Following the recovery period, starting at ZT 9 a.m. (Lights ON), continuous baseline EEG and EMG were recorded for 23 hours from freely moving mice (12 hours lights ON and 11 hours lights OFF), with the last hour of the day reserved for animal care and technical maintenance. Each mouse was connected to a tethered recording system including a headstage/recording cable (1x amplification, custom-made, npi electronics GmbH, Germany) along with a commutator (model SL-20, Dragonfly R&D Inc., USA), mounted on a weight-neutral swivel system (custom-made, Streicher M., Innsbruck, Austria), allowing unrestricted movement of the animal within the recording cage. EEG and EMG signals were independently amplified at 1000x (DPA-2FL Differential Amplifier, npi electronics, Germany). Signals were band-pass filtered between 0.1 and 100 Hz and sampled at 250 Hz (Power1401-3A, Cambridge Electronic Design Ltd., UK). All recording channels included a notch filter at 50 Hz. The raw data were imported into MATLAB - R2024a (MathWorks, USA) for pre-processing and analysis.

### Sleep Scoring

A LABVIEW-based (National Instruments, Austin, TX, USA), semi-automated sleep scoring software^18^ was used. Signals from the left occipital cortex EEG, together with the corresponding EMG, were segmented into 4-second, non-overlapping epochs. Signals from the left occipital cortex EEG, together with the corresponding EMG, were segmented into 4-second, non-overlapping epochs. The software down-sampled the signals to 125 Hz and utilized a decision-tree algorithm based on amplitude-based thresholds for root-mean-square EMG activity and calculated EEG parameters (delta and theta ratios^16^). Each epoch was assigned to one of three vigilance states: WAKE, NREMS, and REMS. The semi-automated sleep scores were manually reviewed and rescored by an experienced scorer if needed to ensure accuracy.

### Epoch selection and annotation of intermediate states

Transition windows were identified from the hypnogram after manual revision. To ensure that the transition was stable enough for experts and to avoid confusion in scoring fluctuating transitions, a custom MATLAB script scanned to detect sequences of 7 consecutive 4-s epochs of one vigilance state (e.g., WAKE) immediately followed by 7 consecutive 4-s epochs of a different state (e.g., NREMS). This procedure ensured that only stable, well-defined transitions were included, while brief or unstable fluctuations in vigilance state were excluded. Each detected window, therefore, consisted of 14 epochs (56 s) representing a period in which a transition occurred. Mice that did not have at least 5 WAKE to NREMS, 5 NREMS to WAKE, 2 NREMS to REMS transitions, and 2 REMS to NREMS transition windows were excluded from further analysis.

To determine the precise timing and duration of these transitions, a separate group of eight trained experts (independent of the original scorer who verified the semi-automated hypnograms) annotated all identified transition windows using a custom MATLAB graphical user interface (Supplementary Fig. 1). To avoid anchoring effects, each 56-s window was randomly shifted by up to ±16 s before presentation Experts were asked to determine whether there is a transition, and if it is a sharp (<1 second) transition. If they believed there was a sharp transition, they could click on the screen (on the EEG, spectrogram, or EMG) at the moment of transition (single 1-second bin) (Supplementary Fig. 1). Regardless of the sharpness, all experts were then invited to select a broader transition period (≤10 seconds), if applicable. EEG spectrograms were limited to 0–40 Hz to facilitate state identification, and all plots were synchronized with a shared time axis for intuitive navigation.

All expert annotations were saved and subsequently used to validate model predictions and quantify inter-expert agreement on the location and boundaries of vigilance-state transitions.

### Inter-rater agreement and characterization of intermediate states

To quantify the consistency among experts in marking vigilance transitions, we computed multiple inter-rater agreement metrics for each event. Expert annotations—either as single time points or as transition intervals—were discretized into 56 one-second bins per transition window, with bins labeled as #1 (state before transition), #2 (during transition), or #3 (state after transition). These annotations were used to calculate pairwise inter-rater agreement in the form of precision, recall, and F1 scores between all expert pairs, and were averaged across transition types. Additionally, we assessed inter-rater reliability using a leave-one-expert-out range criterion: for each transition, an expert’s midpoint was counted as agreeing if it lay within the min–max interval defined by the remaining experts’ midpoints. The per-event reliability score was the proportion of experts meeting this criterion.

To characterize the shape of each transition, we computed transition slope, midpoint, and duration for each expert and each transition window and then averaged it based on transition type. The slope (defined as the steepness of the transition trajectory) was mathematically calculated as the change in hypnodensity probability of the post-transition state over the duration of the transition interval (ΔP/Δt). Consequently, this metric reflects the velocity of the probability shift across the transition trajectory rather than the duration between stable states, and is therefore expressed in arbitrary units.The midpoint was calculated either from the average of the annotated transition interval or directly from the exact time point if no window was selected. Slope was derived from the rate of change in the transition segment, and durations were averaged, excluding zero-duration cases (when one or more experts believed that there was no transition).

### Predicting transition midpoints

To enable the SVM model to automatically detect the transition midpoint with temporal accuracy of one second, we extracted EEG and EMG features in 1-second time bins (125 samples/bin at 125 Hz). From the EEG, absolute band power was calculated across standard frequency bands: Delta (0.1–4 Hz), Theta (4–8 Hz), Alpha (8–13 Hz), Beta (13–30 Hz), and Low Gamma (30–45 Hz). We also calculated complexity and spectral features, including the 1/f spectral slope (25–45 Hz)^19^, Lempel-Ziv complexity (LZc)^20^, and custom DELTA and THETA ratios^18^. Simultaneously, the RMS value of the EMG signal was computed for each 1-second bin. To reduce variability and enhance temporal structure, we applied four smoothing strategies to each feature time series, including Triangular-weighted smoothing, Gaussian smoothing, Prior-2-second averaging and Exponential moving average (EMA). Both smoothed and original features were stored for analysis (see Table 1 for full feature and smoothing details).

**Table 1.**
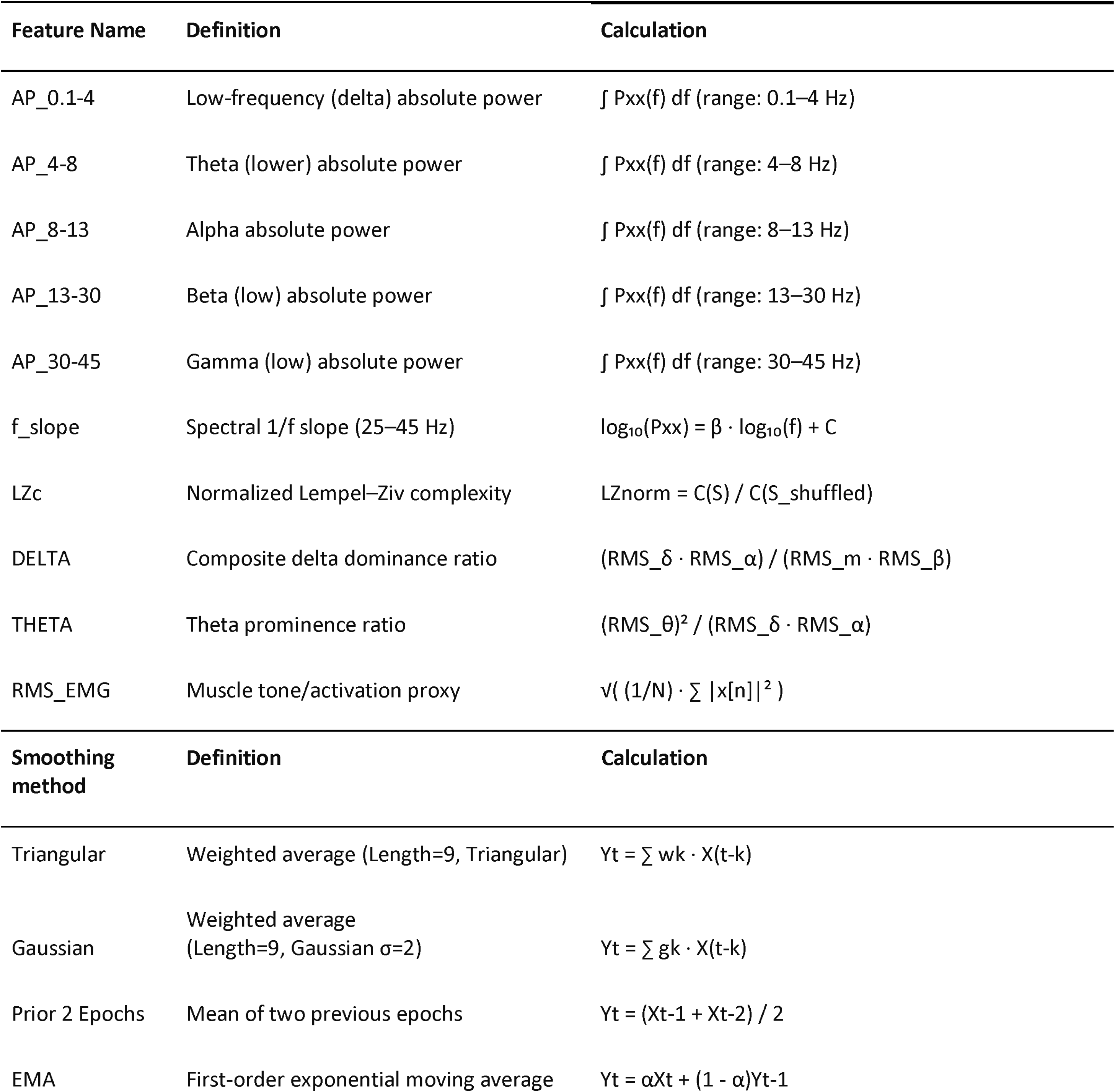
Feature and smoothing details.

| Feature Name | Definition | Calculation |
| --- | --- | --- |
| AP_0.1-4 | Low-frequency (delta) absolute power | $\int P_{xx}(f) df$ (range: 0.1–4 Hz) |
| AP_4-8 | Theta (lower) absolute power | $\int P_{xx}(f) df$ (range: 4–8 Hz) |
| AP_8-13 | Alpha absolute power | $\int P_{xx}(f) df$ (range: 8–13 Hz) |
| AP_13-30 | Beta (low) absolute power | $\int P_{xx}(f) df$ (range: 13–30 Hz) |
| AP_30-45 | Gamma (low) absolute power | $\int P_{xx}(f) df$ (range: 30–45 Hz) |
| f_slope | Spectral 1/f slope (25–45 Hz) | $\log_{10}(P_{xx}) = \beta \cdot \log_{10}(f) + C$ |
| LZc | Normalized Lempel–Ziv complexity | $LZ_{norm} = C(S) / C(S_{shuffled})$ |
| DELTA | Composite delta dominance ratio | $(RMS_{\delta} \cdot RMS_{\alpha}) / (RMS_m \cdot RMS_{\beta})$ |
| THETA | Theta prominence ratio | $(RMS_{\theta})^2 / (RMS_{\delta} \cdot RMS_{\alpha})$ |
| RMS_EMG | Muscle tone/activation proxy | $\sqrt{(1/N) \cdot \sum \vert x[n]\vert ^2}$ |

| Smoothing method | Definition | Calculation |
| --- | --- | --- |
| Triangular | Weighted average (Length=9, Triangular) | $Y_t = \sum w_k \cdot X(t-k)$ |
| Gaussian | Weighted average (Length=9, Gaussian $\sigma=2$) | $Y_t = \sum g_k \cdot X(t-k)$ |
| Prior 2 Epochs | Mean of two previous epochs | $Y_t = (X_{t-1} + X_{t-2}) / 2$ |
| EMA | First-order exponential moving average | $Y_t = \alpha X_t + (1 - \alpha)Y_{t-1}$ |

To build a robust model for identifying transition midpoints, we first filtered the dataset to exclude transitions where fewer than 5 experts agreed on the presence of a transition. We trained SMV classifiers with a linear kernel using data from 6 mice. Classifiers were trained only on feature vectors derived from stable vigilance states, defined as the first 20 seconds (pre-transition) and last 20 seconds (post-transition) of the window, as no transition midpoint was detected in these time windows. Binary classification datasets were constructed for each transition type (e.g., WAKE to NREMS). Model performance in the training dataset was evaluated first using 10-fold cross-validation (*k*=10).

After training on stable segments from 6 mice, the models were applied to full 56-second transition windows from three independent ("unseen") test mice. We first assessed how well the model reproduces the stable states in the three unseen mice. For each 56-s window, the SVM produced a per-second probability p_t_ of the post state. In the first 20 s and last 20 s, we converted probabilities to hard labels using confidence bands: p_t_<0.20 considered as pre-transition and p_t_>0.80 considered as post-transition. A bin of one second was counted accurately when the model label matched the expert majority (for example, if the majority of experts consider the state as pre-transition and the p_t_ is 0.04, it is considered an accurate bin). For each window, we calculated the accuracy as the percentage and summarized mean ± SEM per transition type. Then, we calculated how accurately the model can find the transition midpoint. The model-predicted midpoint was defined as the first time bin where the smoothed posterior probability exceeded 0.5. To address the inherent subjectivity in expert annotations, we evaluated prediction accuracy using a graded agreement metric. For each transition window, we counted how many of the 8 experts’ individual annotation windows (defined as the interval between each expert’s marked start and end time) contained the model-predicted midpoint. We then calculated the percentage of windows where this count met or exceeded a threshold of ≥1, ≥2, ≥3, or ≥4 experts. The same graded metric was applied symmetrically to expert annotations using a leave-one-out approach, where each expert’s midpoint was checked against the remaining 7 experts’ individual windows, providing a human baseline for direct comparison. Model performance was evaluated using leave-3-out cross-validation across all C(9,3) = 84 possible combinations of training (6 mice) and test (3 mice) sets, and results are reported as mean ± SEM across folds.

### Statistical Analysis

Statistical analyses were performed using GraphPad Prism software (version 10.0; GraphPad Software, San Diego, CA). Graphical representations were generated primarily using MATLAB (MathWorks) and GraphPad Prism. Data distribution and normality were assessed using the D’Agostino-Pearson omnibus test and the Shapiro-Wilk test. All statistical tests were two-tailed, and significance was considered at a 95% confidence level (α = 0.05).

For normal distributed data, group comparisons were performed using one-way analysis of variance (ANOVA), followed by Tukey’s multiple comparisons test to correct for multiple testing. For non-normal distributed data, group comparisons were performed using the Kruskal-Wallis test followed by Dunn’s multiple comparisons test. Data are presented as mean ± standard error of the mean (SEM). Non-normal distributed data was represented as median and quantile range. The p-values for all relevant comparisons are reported in the corresponding supplementary tables and figure legends.

## Results

### Distribution of intermediate vigilance states

From the initial pool of 11 animals, transitions were extracted for four vigilance state changes: WAKE to NREMS (W-N), NREMS to WAKE (N-W), NREMS to REMS (N-R), and REMS to NREMS (R-N). Mice M1LJ15, and M1LJ26 did not meet the minimum transition count for either NR or RN and were excluded from further analysis (Supplementary Table 1). In total, 394 transitions across 9 mice were selected for expert annotation (Figure 1), including 145 WAKE to NREMS, 150 NREMS to WAKE, 52 NREMS to REMS, and 47 REMS to NREMS transitions. Two mice (M1LJ15 and M1LJ26) did not meet the minimum transition count criteria for either NREMS to REMS or REMS to NREMS and were excluded from further analysis (Supplementary Table 1).

**Figure 1.**
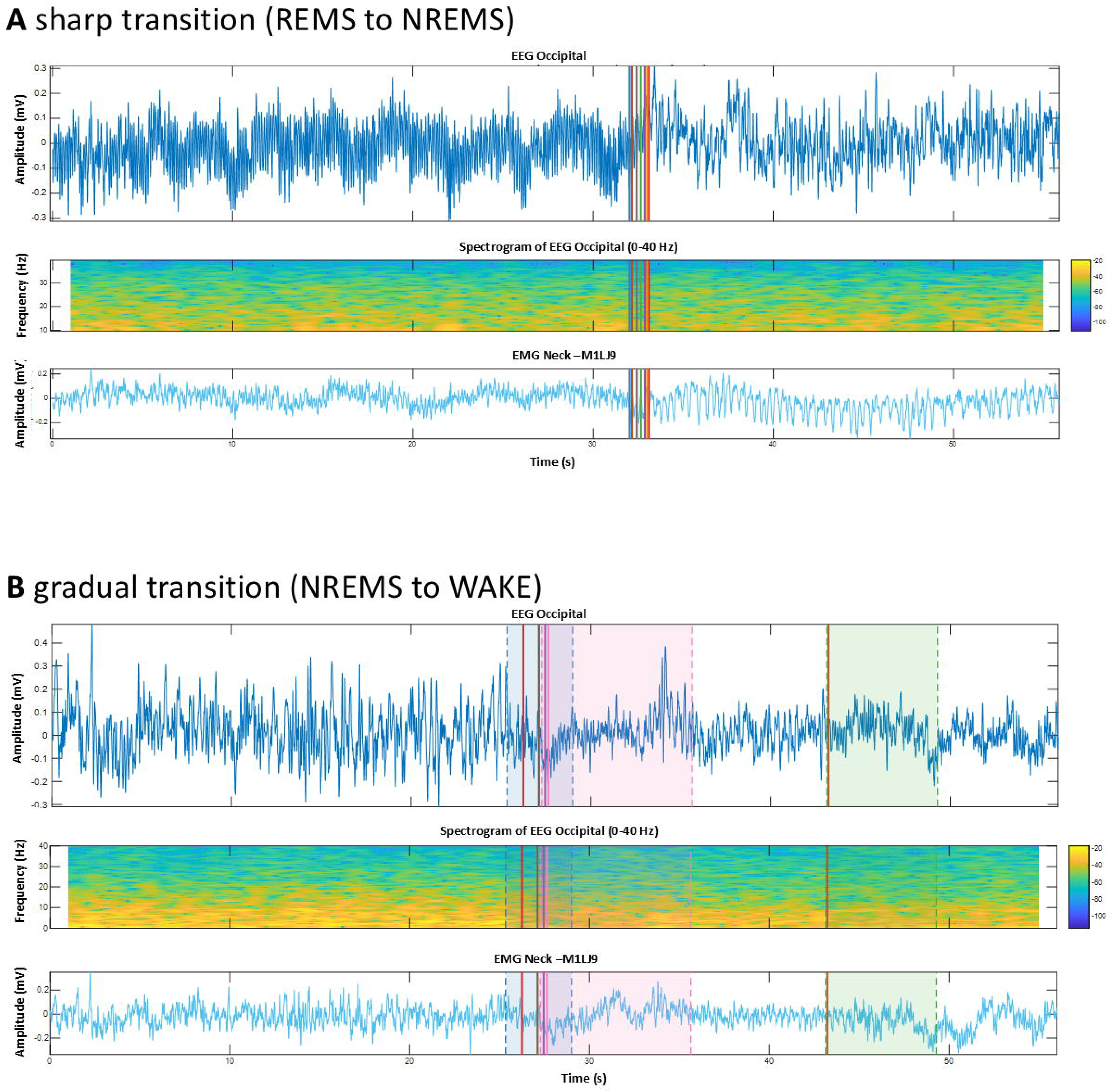
Representative example of (A) sharp and (B) gradual transition based on the annotation of 8 experts. EEG recording (top panel), corresponding spectrogram (middle panel) and simultaneously recorded EMG (bottom panel) recorded in the nuchal muscle. Colored vertical lines indicate an expert-annotated transition midpoint.

### Inter-Rater Agreement Across Experts

The consistency of expert annotations across sleep-wake transitions is visualized in Figure 2 and summarized in Table 2.

**Figure 2.**
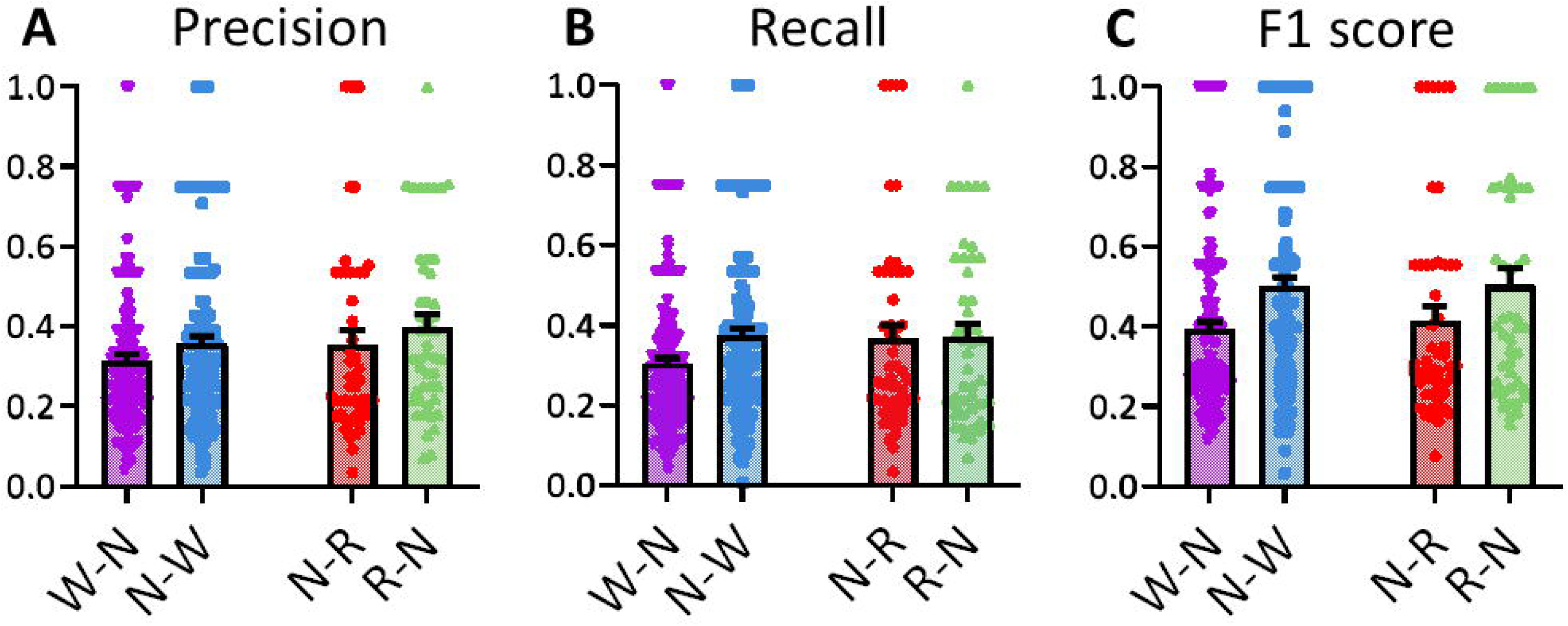
Summary of inter-rater agreement across experts. (A) Precision, (B) Recall and (C) F1 score reflecting bin-level overlap in the temporal labelling of transitions. WAKE to NREMS (W-N), NREMS to WAKE (N-W), NREMS to REMS (N-R), and REMS to NREMS (R-N).

**Table 2.** Summary of inter-rater agreement across experts.

| Measure | WAKE to NREM | NREM to WAKE | NREM to REM | REM to NREM |
| --- | --- | --- | --- | --- |
| Precision | 0.314 ± 0.015 | 0.358 ± 0.017 | 0.355 ± 0.034 | 0.398 ± 0.032 |
| Recall | 0.304 ± 0.014 | 0.374 ± 0.017 | 0.367 ± 0.034 | 0.370 ± 0.032 |
| F1 Score | 0.394 ± 0.018 | 0.502 ± 0.022 | 0.416 ± 0.036 | 0.505 ± 0.041 |
| inter-rater reliability | 0.653 ± 0.049 | 0.688 ± 0.077 | 0.617 ± 0.018 | 0.652 ± 0.021 |

F1 scores, reflecting bin-level overlap in the temporal labeling of transitions, were low to moderate and showed greater variability across transitions. Highest F1 scores were found for REMS to NREMS (0.505 ± 0.041) and NREMS to WAKE (0.502 ± 0.022), whereas WAKE to NREMS (0.394 ± 0.018) and NREMS to REMS (0.416 ± 0.036) had comparatively lower scores, reflecting increased variability in temporal annotation for these transitions.

To further quantify temporal agreement, we calculated inter-rater reliability. This measure captures consistency in identifying the exact time of transition onset and shows overall moderate values. Inter-rater reliability was highest for NREMS to WAKE (0.688 ± 0.077) and REMS to NEREMS (0.652 ± 0.021), with WAKE to NREMS (0.653 ± 0.049) and NREMS to REMS (0.617 ± 0.018) showing slightly lower values.

### Transition Dynamics

We next quantified the temporal characteristics of expert-annotated vigilance state transitions by analyzing their slope, duration, and midpoint variability (visualized in Figure 3 and summarized in Table 3).

**Figure 3.**
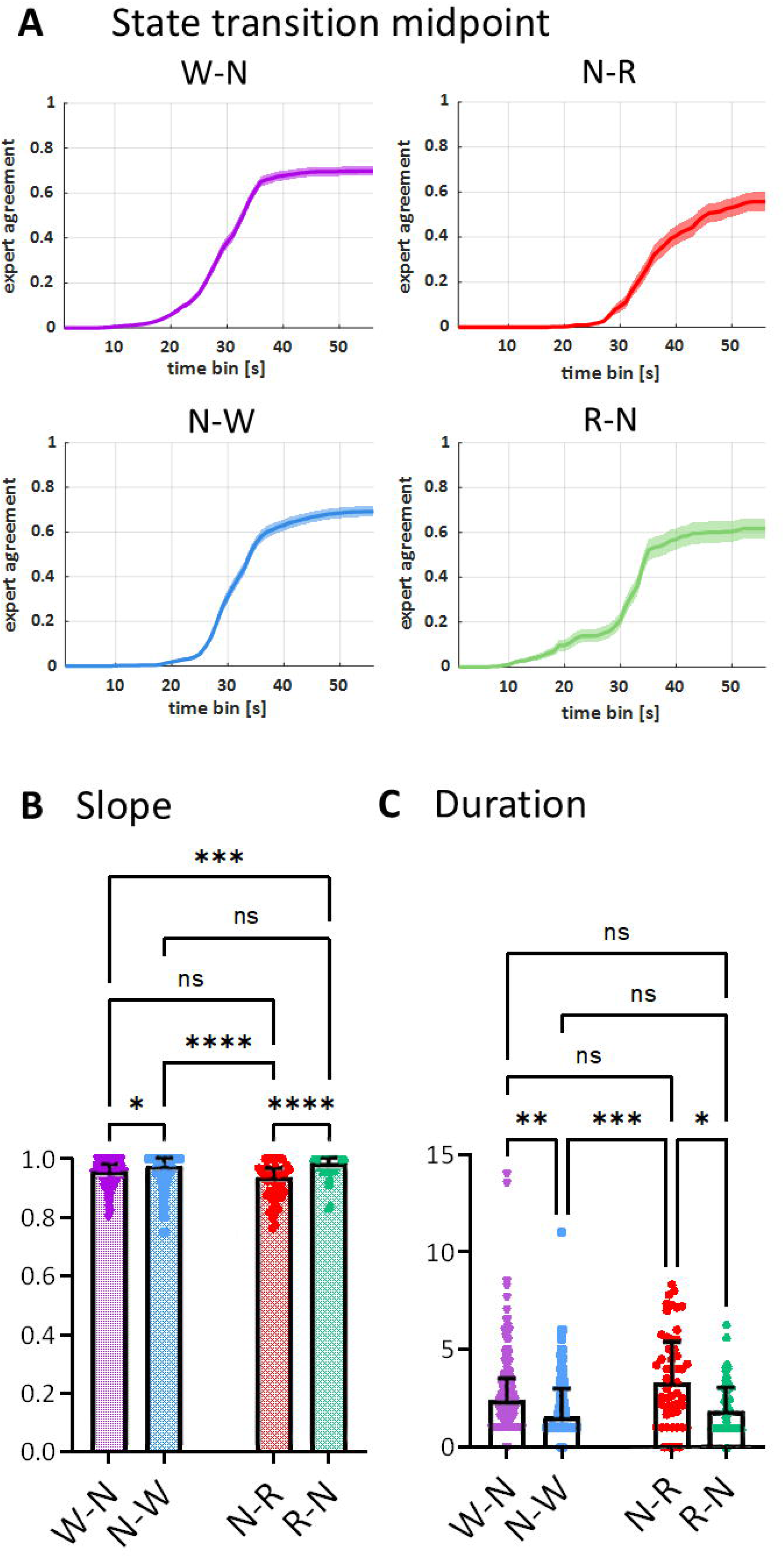
Temporal characteristics of expert-annotated vigilance state transitions. (A) Probability of entering post-transition state based on scoring provided by 8 experts (B) slope of annotated transitions. W-N to N-W: < 0.05, W-N to R-N: < 0.0001, N-W to N-R: p<0.0001, N-R to R-N: < 0.0001; according to Kruskal-Wallis test followed by Dunn’s multiple comparison test (C) duration of annotated transitions. W-N to N-W: < 0.01; N-W to N-R: < 0.001; N-R to R-N: p < 0.01; according to Kruskal-Wallis test followed by Dunn’s multiple comparison test WAKE to NREMS (W-N), NREMS to WAKE (N-W), NREMS to REMS (N-R), and REMS to NREMS (R-N). In (A), lines represent the mean, shaded areas indicate ±SEM. (B, C) Each bar represents the median, whiskers represent quantiles.

**Table 3.** Temporal characteristics of expert-annotated vigilance state transitions.

| Measure | WAKE to NREM | NREM to WAKE | NREM to REM | REM to NREM |
| --- | --- | --- | --- | --- |
| Slope (a.u.) | $0.951 \pm 0.003$ | $0.962 \pm 0.004$ | $0.920 \pm 0.010$ | $0.975 \pm 0.005$ |
| Duration (s) | $2.802 \pm 0.170$ | $2.127 \pm 0.125$ | $3.570 \pm 0.325$ | $2.133 \pm 0.201$ |
| Midpoint range (s) | $9.150 \pm 0.620$ | $7.498 \pm 0.630$ | $11.242 \pm 1.020$ | $9.076 \pm 1.515$ |

The slope was highest for REMS to NREMS transitions (0.975 ± 0.005) and lowest for NREMS to REMS transitions (0.920 ± 0.010), with intermediate values for NREMS to WAKE (0.962 ± 0.004) and WAKE to NREMS (0.951 ± 0.003). Kruskal-Wallis test followed by Dunn’s multiple comparison test revealed that the slope of NREMS to REMS transitions was significantly shallower than NREMS to WAKE (*p* < 0.0001), and REMS to NREMS (*p* < 0.0001) transitions.

Duration, defined as the time between the start and end of the transition as annotated by experts (seconds), varied significantly across transition types (Table 3, Fig. 2). The longest transitions were observed for NREMS to REMS (3.570 ± 0.325 s), followed by WAKE to NREMS (2.802 ± 0.170 s), REMS to NREMS (2.133 ± 0.201 s), and NREMS to WAKE (2.127 ± 0.125 s). Kruskal-Wallis test with Dunn’s post hoc test indicated that NREMS to REMS transitions were significantly longer than all other transition types (all *p* < 0.05 and lower), while WAKE to NREMS is significantly longer than NREMS to WAKE (p < 0.01).

### Predicting midpoints of transition windows using SVM

After excluding transition windows that did not meet the minimum requirement of at least five expert annotations (see Methods), a total of 118 WAKE to NREMS, 116 NREMS to WAKE, 38 NREMS to REMS, and 33 REMS to NREMS transitions were included in the analysis (Supplementary Table 2).

SVM models were trained separately for each transition type using only the first and last 20 time bins (stable states) of each transition window for six mice. Confusion matrices showed high classification performance during training, with accuracies close to 99–100% across all transition types (Supplementary Figure 2). Then, in three randomly selected “unseen” mice, the model was used to predict the probability of entering the second state (e.g., NREMS in a WAKE to NREMS transition) across all 56 bins (Figure 4). We first assessed how well the model reproduces the stable states in the three unseen mice, which yielded the accuracies of 88.67 ± 1.44 %, 90.45 ± 1.10%, 81.05± 3.23%, and 86.89 ± 4.31% for WAKE to NREMS, NREMS to WAKE, NREMS to REMS, and REMS to NREMS transitions, respectively (Table 4, Figure 5). To evaluate the SVM model’s ability to localise the transition midpoint, we applied a graded metric in which the model-predicted midpoint was required to fall within the annotation window of at least *k* experts (*k* = 1, 2, 3, 4), and compared model performance against a human baseline computed using the same leave-one-out criterion across all 84 cross-validation folds (Figure 4, Table 4). Under the ≥1 expert criterion, model accuracy was 50.2 ± 1.4% for WAKE to NREMS, 40.0 ± 0.7% for NREMS to WAKE, 52.7 ± 1.8% for NREMS to REMS, and 46.9 ± 1.6% for REMS to NREMS transitions. These values were broadly comparable to the human baseline under the same criterion (49.6 ± 4.5%, 45.7 ± 2.6%, 74.3 ± 3.6%, and 45.5 ± 7.2% respectively), with the exception of NREMS to REMS where the model underperformed experts at the ≥1 threshold but converged toward expert-level performance at stricter thresholds (≥2: model 29.9 ± 1.7% vs expert 33.9 ± 5.2%). Accuracy declined monotonically as the threshold increased for both model and experts, as at ≥4 experts, both model and expert agreement approached zero for most transition types, consistent with the low inter-rater reliability reported above. (Figure 6).

**Figure 4.**
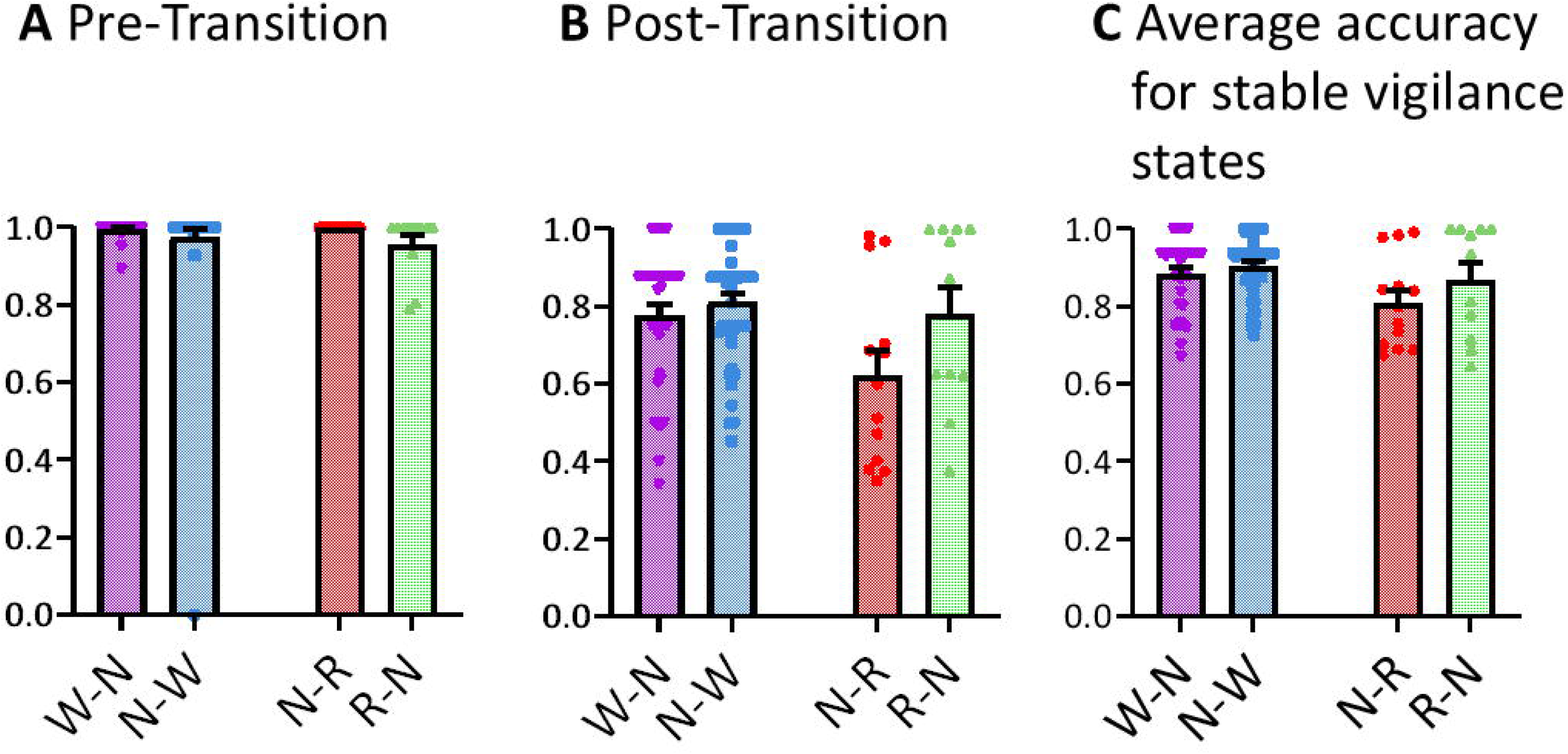
How accurate SVM can detect stable vigilance state in comparison to experts annotations. (A) 20 s time window pre-transition, (B) 20 s time window post-transition and (C) the average accuracy of the model in predicting stable vigilance states. WAKE to NREMS (W-N), NREMS to WAKE (N-W), NREMS to REMS (N-R), and REMS to NREMS (R-N).

**Figure 5:**
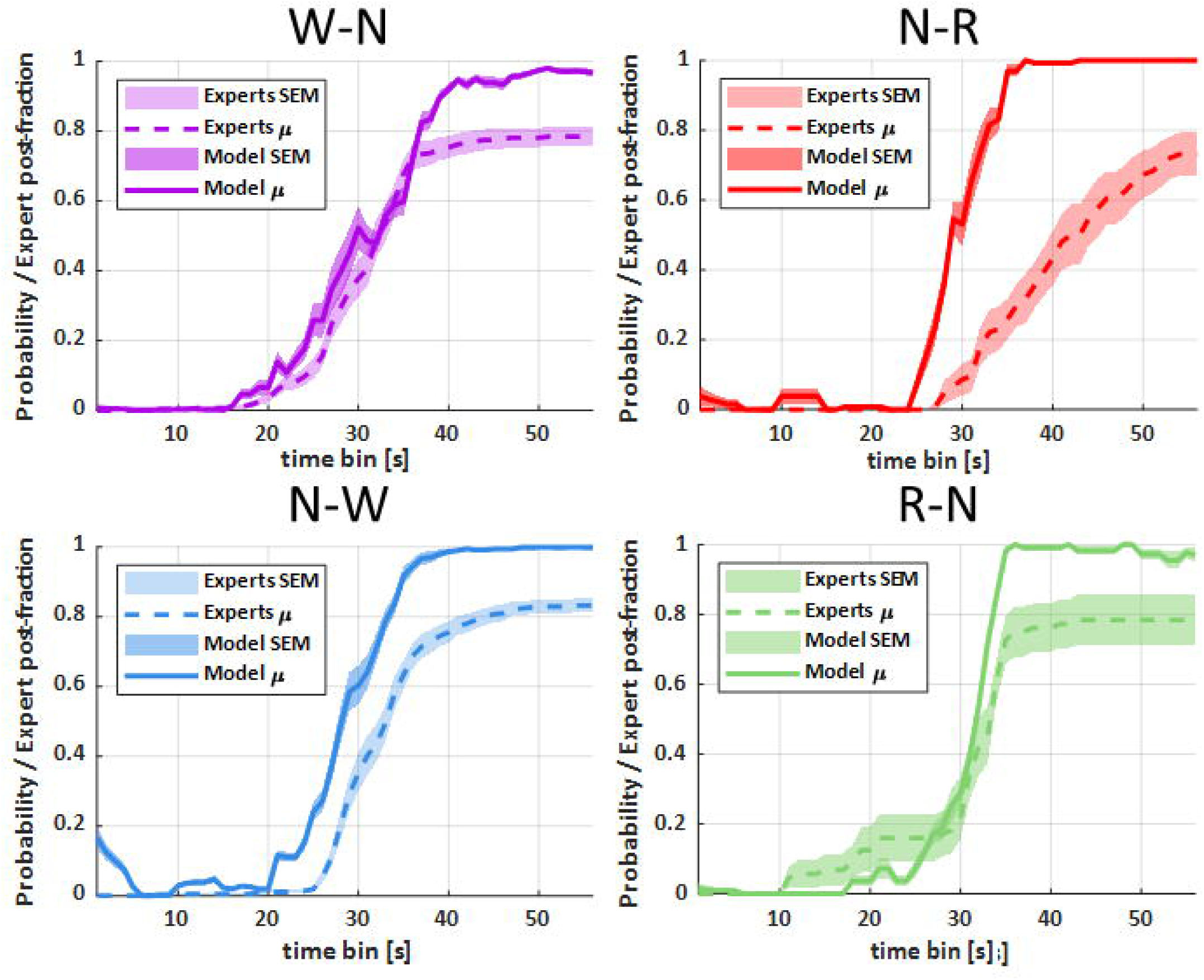
Comparison between experts annotated and SVM predicted probability of entering post-transition state. WAKE to NREMS (W-N), NREMS to WAKE (N-W), NREMS to REMS (N-R), and REMS to NREMS (R-N). Lines represent mean, shaded areas indicate ±SEM.

**Figure 6:**
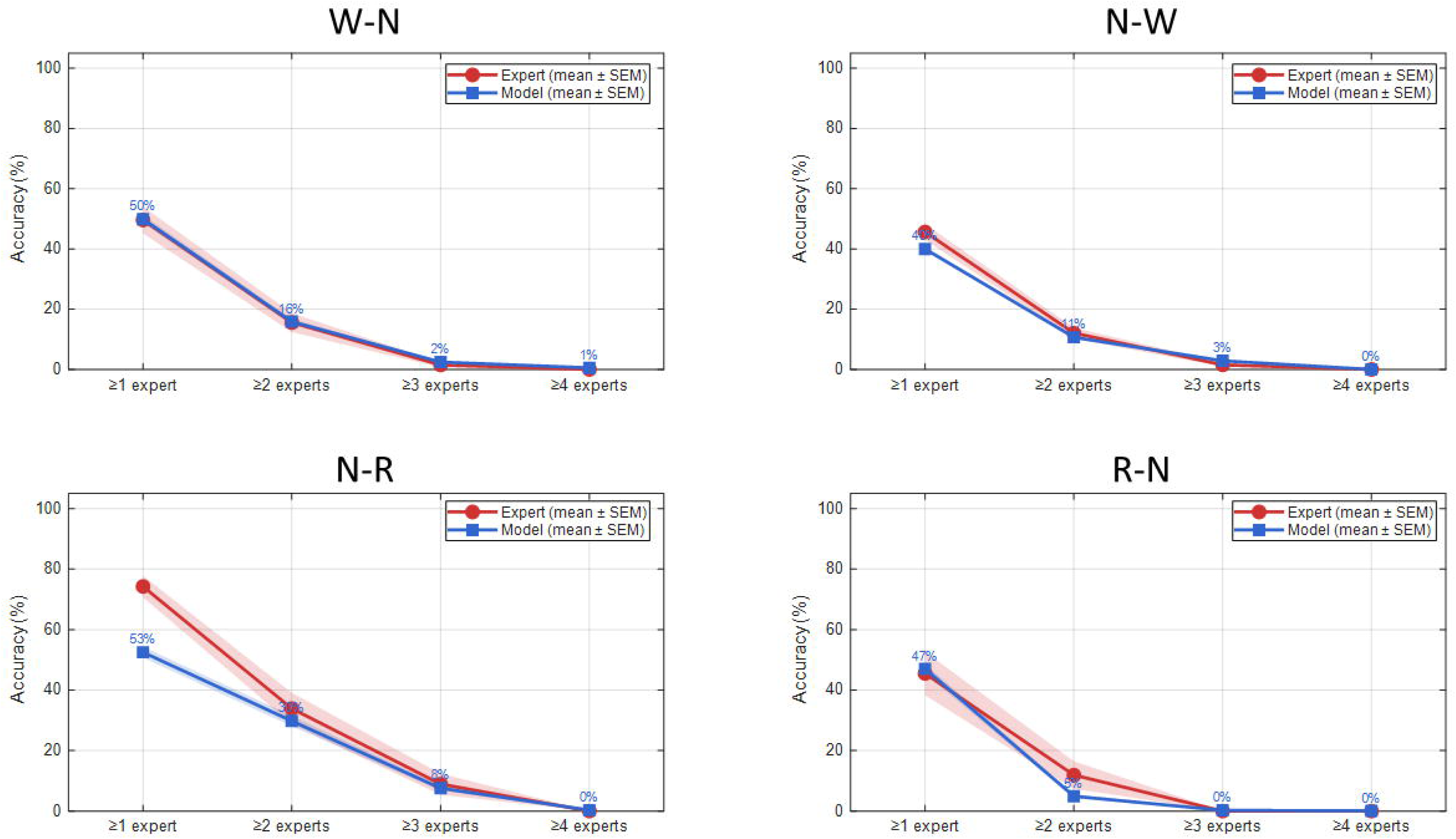
Model-graded agreement compared to expert-graded agreement across different thresholds and transition types. For each transition window, how many of the 8 experts’ individual annotation windows contained the model-predicted midpoint were counted. The percentage of windows where this count met or exceeded a threshold of ≥1, ≥2, ≥3, or ≥4 experts was then calculated. The same graded metric was applied symmetrically to expert annotations using a leave-one-out approach, where each expert’s midpoint was checked against the remaining 7 experts’ individual windows. WAKE to NREMS (W-N, top-left), NREMS to WAKE (N-W, top-right), NREMS to REMS (N-R, bottom-left), and REMS to NREMS (R-N, bottom-right). Lines represent mean, shaded areas indicate ±SEM.

**Table 4.**
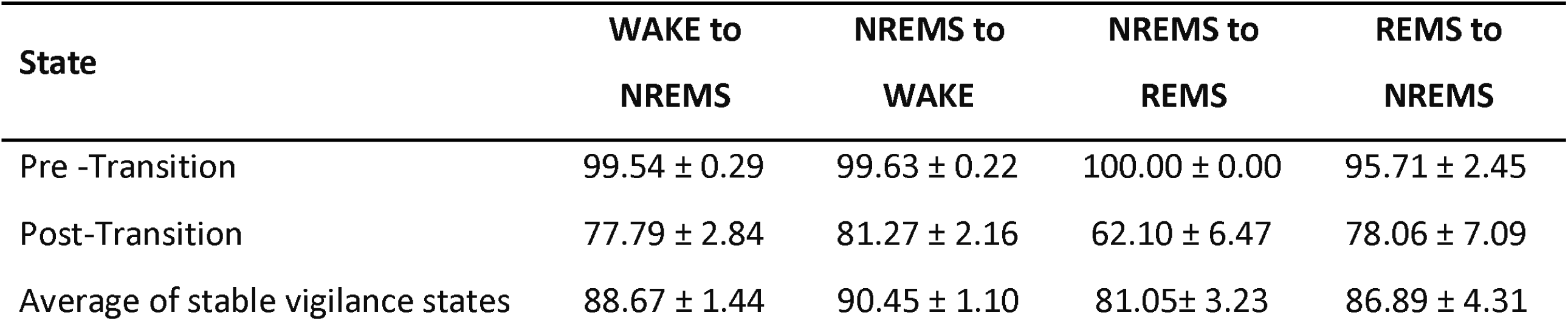
Accuracy of SVM in predicting stable vigilance states (mean ± SEM)

## Discussion

In this study, we characterized the temporal dynamics of sleep–wake transitions in mice and demonstrated that intermediate vigilance states can be annotated and predicted using quantitative EEG/EMG features. By combining expert-based labeling with machine learning, our approach provides a reproducible framework for quantifying the temporal midpoint of sleep-wake transitions and for characterizing their associated dynamics — including duration, slope, and inter-expert variability — properties that are complementary to, rather than replacements for, conventional epoch-based sleep scoring. Our primary findings demonstrate three key observations. First, human experts exhibited significant variability and moderate-to-low agreement in defining transition midpoints, particularly for NREMS to REMS transitions. Second, different scored transition types possess distinct temporal signatures, with NREMS to REMS transitions being significantly slower and longer than all others. Third, stable-state EEG/EMG features were sufficient for machine-learning models to detect transition timing with acceptable accuracy, despite the variability observed in human annotations.

The pursuit of high-temporal resolution in transition detection addresses a fundamental divergence in sleep literature: while some models suggest a sharp bifurcation point^3,21^, others describe transitions as progressive cycles lasting tens of seconds^22^. By establishing an objective framework for identifying transition midpoints, our method moves beyond the ’as-if’ assumption of discrete boundaries. This granularity is essential for future studies seeking to correlate sudden shifts in dynamical regimes with specific cognitive deficits or the paroxysmal activity seen in neurological conditions like epilepsy, where the transition period itself may serve as a window of heightened vulnerability.

In total, 394 transitions from 9 mice were analyzed for this study (see supplementary table 1). This low number in transitions can be explained by our strict stability criterion; only transitions flanked by 7 consecutive epochs of each state were included in the analysis. This leads to the exclusion of many brief NREMS and REMS bouts.

A subtle but consistent finding in our data was an asymmetry in the duration of state transitions. Specifically, WAKE to NREMS was longer than NREMS to WAKE, while the converse was true for transitions between NREMS to REMS and REMS to NREMS. This difference is small (on the order of a few seconds, also seen by Sánchez-López et al.^23^) and must be interpreted cautiously, given the variance. Methodologically, standard sleep scoring requires a minimum duration (commonly a 4-second epoch^24^) to confidently define a state, making boundaries inherently fuzzy. For example, defining the precise end of NREMS (or beginning of REMS) often relies on identifying the “last spindle”^14^, which may require a retrospective window, artificially elongating the perceived transition out of NREMS. Further, between NREMS and REMS there is often a “pre-REM”^6^ that differs slightly in electrophysiological profile from both REMS and NREMS but is nonetheless quite short, potentially complicating scoring of the transition^22,25^. Similarly, the transition from WAKE into NREMS found in the present study may pass through a stage similar to NREM1 in humans, although less pronounced^14^. Together, these points suggest that the transition-duration asymmetry may arise from classification challenges due to intermediate states and NREMS heterogeneity, making the precise identification of start and end points inherently uncertain. Future studies are needed to map this out in more detail.

It is worth mentioning that, although the results here and the existence of intermediary states may suggest a gradual transition between sleep states, transitions may also be abrupt. For example, Li et al. 2025^21^ in humans, showed the existence of a bifurcation point, signaling the irrevocable transition from one state to the next, even though the neural dynamics themselves may change gradually. On the other hand, there may be physiological reasons for transition-duration asymmetry. Interestingly, simultaneous single-unit recordings of nucleus accumbens and locus coeruleus in mice show a similar disparity in the two transitions, as the NREMS to WAKE transition is clearly more abrupt than the WAKE to NREMS transition^26^. This is also comparable to the observation in humans^27^.

The exceptional length and variability of NREMS to REMS transitions suggest that REMS entry is not triggered by a discrete event but requires the gradual accumulation of permissive conditions, most likely the slow withdrawal of the inhibitory tone that gates REMS onset. This is highly consistent with single-unit and calcium imaging recordings in mice showing that REM-off neurons in the ventrolateral periaqueductal grey decrease their firing rate gradually across NREMS bouts, reaching a minimum just before REMS onset, and abruptly increasing at REMS termination^28^. The long NREMS to REMS transition duration we observe may be the EEG-level correlate of that disinhibition build-up. Conversely, the comparatively sharp REMS to NREMS boundary suggests that REMS termination is an active, rapid process driven by the quick re-engagement of these REM-off populations^28^. Similarly, the asymmetry between WAKE to NREMS and NREMS to WAKE transitions suggests these are mechanistically distinct events rather than mirror images of the same switch. Sleep onset requires the gradual build-up of homeostatic sleep pressure and inhibitory tone sufficient to overcome tonic arousal drive, whereas waking can be triggered rapidly by a single strong arousal signal.

While no study directly compares the temporal dynamic of NREMS to REMS and REMS to NREMS transitions, several existing single-unit activity studies indicate that the NREMS to REMS transition is gradual and can take several seconds^29^. Ultimately, the present data were not designed to answer this question. A dedicated analysis tracking multiple markers (e.g., subcortical measurements of sleep-regulating regions, etc.) at high temporal resolution would be required to explore the role of NREMS and REMS substrates in shaping these transition dynamics.

Despite standardized instructions and experienced scorers, inter-rater reliability across experts remained moderate to low. The low F1 scores and moderate reliability for WAKE to NREMS and NREMS to REMS transitions indicate that pinpointing the exact onset of a new vigilance state is a challenging task. This ambiguity was most evident for the NREMS to REMS transition, which was consistently annotated as the longest and slowest (i.e., the shallowest slope). As highlighted above, the low F1 score for NREMS to REMS may suggest that the perceived “slowness” could stem from the absence of a single, universally used biomarker for NREMS and REMS sleep onset, leading experts to rely on differing internal criteria, like theta to delta ratio or EMG amplitude. The moderate to low agreement for different types of transitions emphasizes the need for new analytical frameworks that acknowledge the continuous nature of vigilance, rather than enforcing arbitrary boundaries through epoch-based scoring.

We purposefully avoided calculating the transition duration directly from the SVM for methodological reasons. In many machine learning approaches, the output—which is often the distance to a decision boundary fed into a logistic function—is mathematically compressed toward extreme probabilities (approaching 0 or 1). This process artificially sharpens the predicted transition, yielding durations that may not reflect the true underlying biological dynamics. A similar problem arises when models are trained only on fixed-sized epochs of WAKE and NREMS and then applied to find transition states between them^9^. To mitigate these artifacts, we instead relied on eight expert scorers to manually define the transition boundaries within a larger window, and then used the machine learning approach solely to determine the midpoint. Despite the "noise" of human subjectivity, our SVM model demonstrated that EEG and EMG features within the transition window contain sufficient information to predict the expert-defined midpoint. The model’s ability to achieve 63-85% accuracy in matching the range of expert annotations validates the idea that a consistent, albeit complex, signal exists within the data. This provides support for the development of automated, objective systems for sleep scoring, which could resolve the inter-rater reliability issues that have long plagued the field.

## Limitations

A potential limitation of this study is our primary reliance on occipital EEG derivations. While previous work has demonstrated that occipital signals provide superior performance for the automated discrimination of WAKE and REMS in mice ^1,3^, this spatial focus may not capture the full regional heterogeneity of sleep-state transitions. Specifically, a fronto-parietal configuration might better resolve the distinct onset of frontal delta power during NREMS or parietal theta during REMS. Given that vigilance-state transitions can occur asynchronously across the cortex, the temporal dynamics and midpoints identified here should be interpreted as specific to the occipital region; further research is required to determine the extent of spatiotemporal ’lag’ across the rostro-caudal axis during these transitions.

Only male C57BL/6N mice were used in this study to minimize between-animal variability in sleep architecture. Oestrous cycle-related fluctuations in sleep in female mice are well-documented ^30^ and would introduce an additional source of variability into expert annotation, which could conflate biological variation with scorer disagreement.

Another potential limitation of the current study is its reliance solely on EEG and EMG recordings, which led to low F1 scores and high variability in epoch-level overlap during vigilance-state transitions. Because state boundaries can be gradual and ambiguous, our ground-truth data regarding these transitions should be interpreted with caution. Future work incorporating additional physiological and behavioural metrics, such as respiration^31^, single-unit recordings from key sleep-wake regulatory centres (e.g., LC and VLPO)^29^, and video tracking, is needed to delineate transition periods more robustly.

Although EEG and EMG features captured transition dynamics effectively, additional modalities (e.g., Local field potentials, respiration, pupil diameter) may refine the definition of transition onset. Another limitation is the our sample size for NREMS to REMS (n=38) and REMS to NREMS (n=33) transitions was relatively small, a consequence of the inherent difficulty in capturing these events and the necessary exclusion of two mice. On the other hand, we could not detect enough number of stable REMS to WAKE transitions to be included in the study.

The SVM model was trained using the expert-annotated midpoints as its "ground truth." It is therefore "learning" the central tendency of a subjective process. While it provides more consistency, it is still biased by the same human-defined criteria. Future studies using unsupervised clustering to pinpoint transition states are necessary.

## Conclusions

This study demonstrates that intermediate vigilance states in rodents are inherently ambiguous and annotated with only moderate consistency across experts. Transition dynamics differ markedly by transition type, with REMS entry being the most gradual and REMS exit the most abrupt. Despite this variability, simple machine-learning classifiers trained on stable states can reliably infer transition midpoints. These findings highlight the potential of data-driven approaches to complement or replace manual scoring and support a more continuous, mechanistic view of vigilance state regulation.

## Acknowledgments

We would like to extend our appreciation to Yasin Aküyz, Snezana Djordjevic, Mona Knoblauch, Catharina Hedenig, Matilde Lupi, and Alp Altunkaya, all from TUM University Hospital, School of Medicine and Health, Clinic of Anesthesiology and Intensive Care, Technical University of Munich, Germany, for their collaborative work in scoring the transition state and for the fruitful discussions.

## Disclosure statement

Financial disclosure: The authors declare that they have no financial arrangements or connections that could be perceived as influencing the work reported in this article.

Non-financial disclosure: The authors declare that they have no non-financial conflicts of interest related to the work reported in this article.

## Data Availability Statement

The data supporting the findings of this study are available from the first author or the corresponding author upon reasonable request.

---

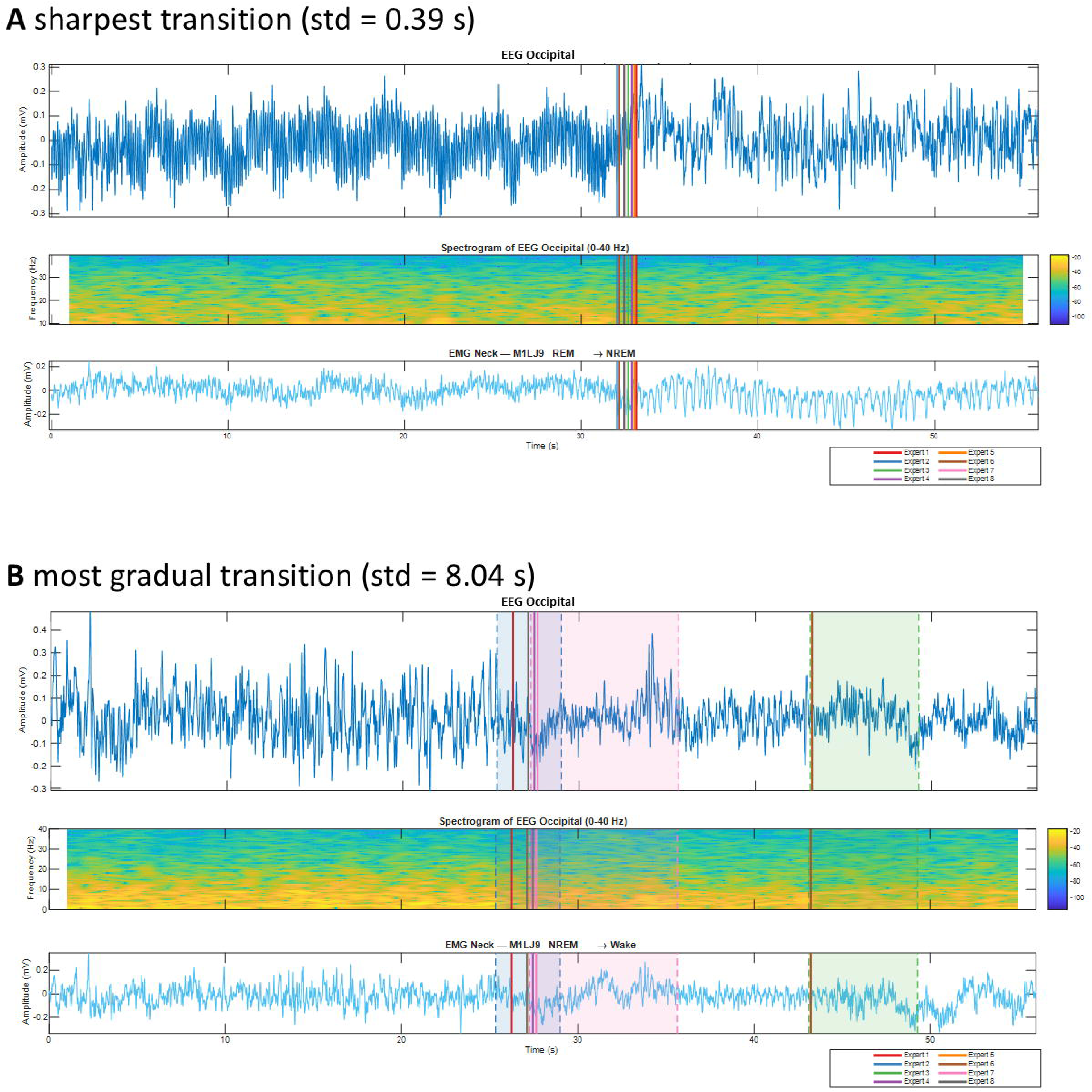

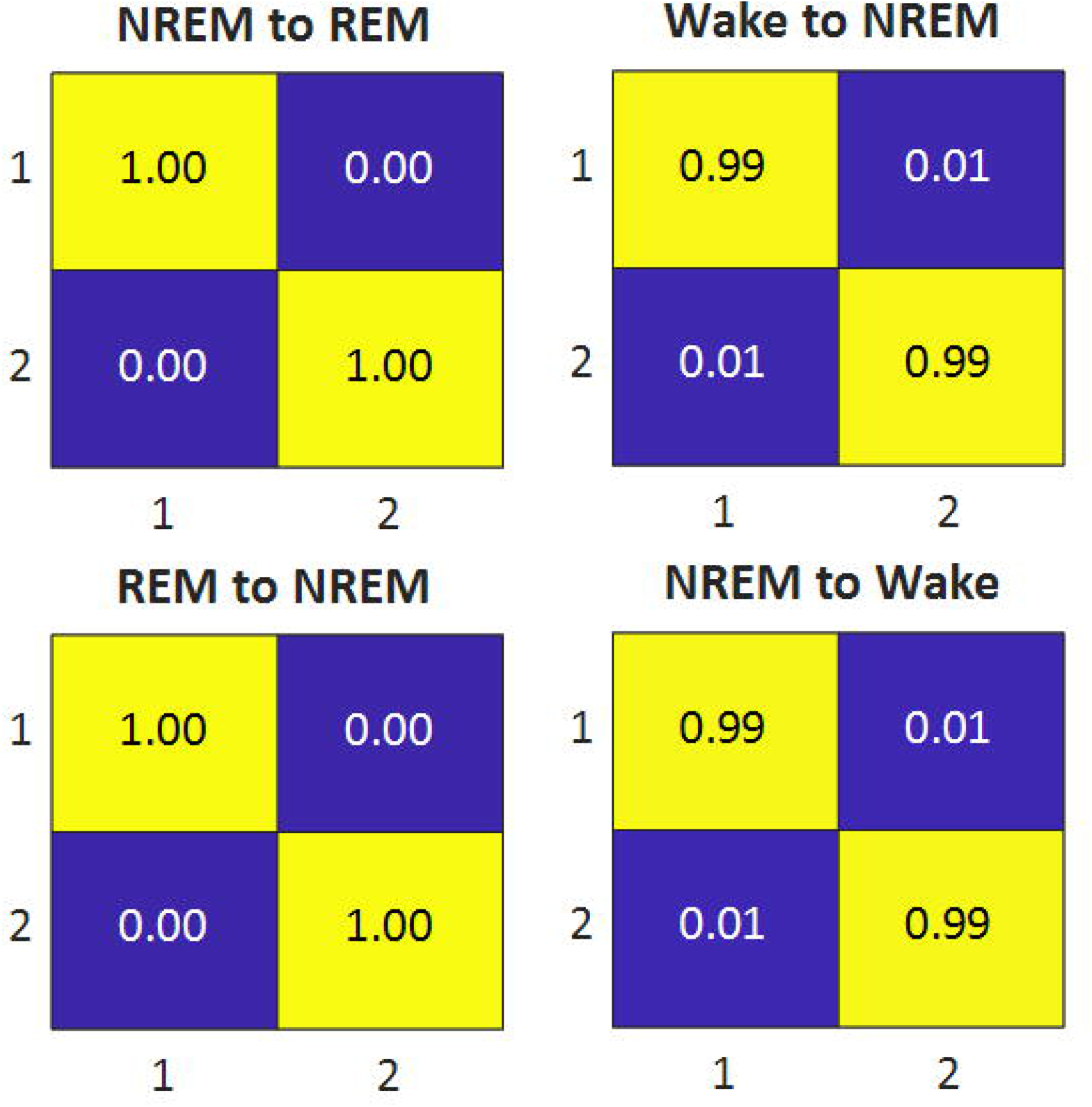

## Notes

### Competing Interest Statement

The authors have declared no competing interest.

