## Supplementary Data for "Characterizing Transition State in Mouse Vigilance with EEG–EMG Hypnodensity"

\# shared last authors

**Supplementary Table 1.** Summary of vigilance state transitions	(shaded rows indicate excluded animals)

| **Mouse** | **WAKE to NREMS** | **NREMS to WAKE** | **NREMS to REMS** | **REMS to NREMS** |
| --- | --- | --- | --- | --- |
| **M1LJ10** | 18 | 19 | 6 | 6 |
| **M1LJ11** | 15 | 15 | 6 | 6 |
| **M1LJ13** | 10 | 10 | 6 | 6 |
| **M1LJ15** | 6 | 6 | 0 | 0 |
| **M1LJ16** | 16 | 17 | 4 | 3 |
| **M1LJ17** | 15 | 16 | 6 | 6 |
| **M1LJ18** | 15 | 15 | 6 | 6 |
| **M1LJ26** | 6 | 5 | 0 | 0 |
| **M1LJ29** | 17 | 15 | 6 | 6 |
| **M1LJ30** | 17 | 15 | 6 | 2 |
| **M1LJ9** | 22 | 28 | 6 | 6 |
| **total** | 145 | 150 | 52 | 47 |

**Supplementary Table 2.** Transition windows, which are excluded from training session (due to lack of consensus between experts)

|  | **WAKE to NREMS** | **NREMS to WAKES** | **NREMS to REMS** | **REMS to NREMS** |
| --- | --- | --- | --- | --- |
| Total Number | 143 | 145 | 48 | 46 |
| Excluded | 25 | 29 | 10 | 13 |

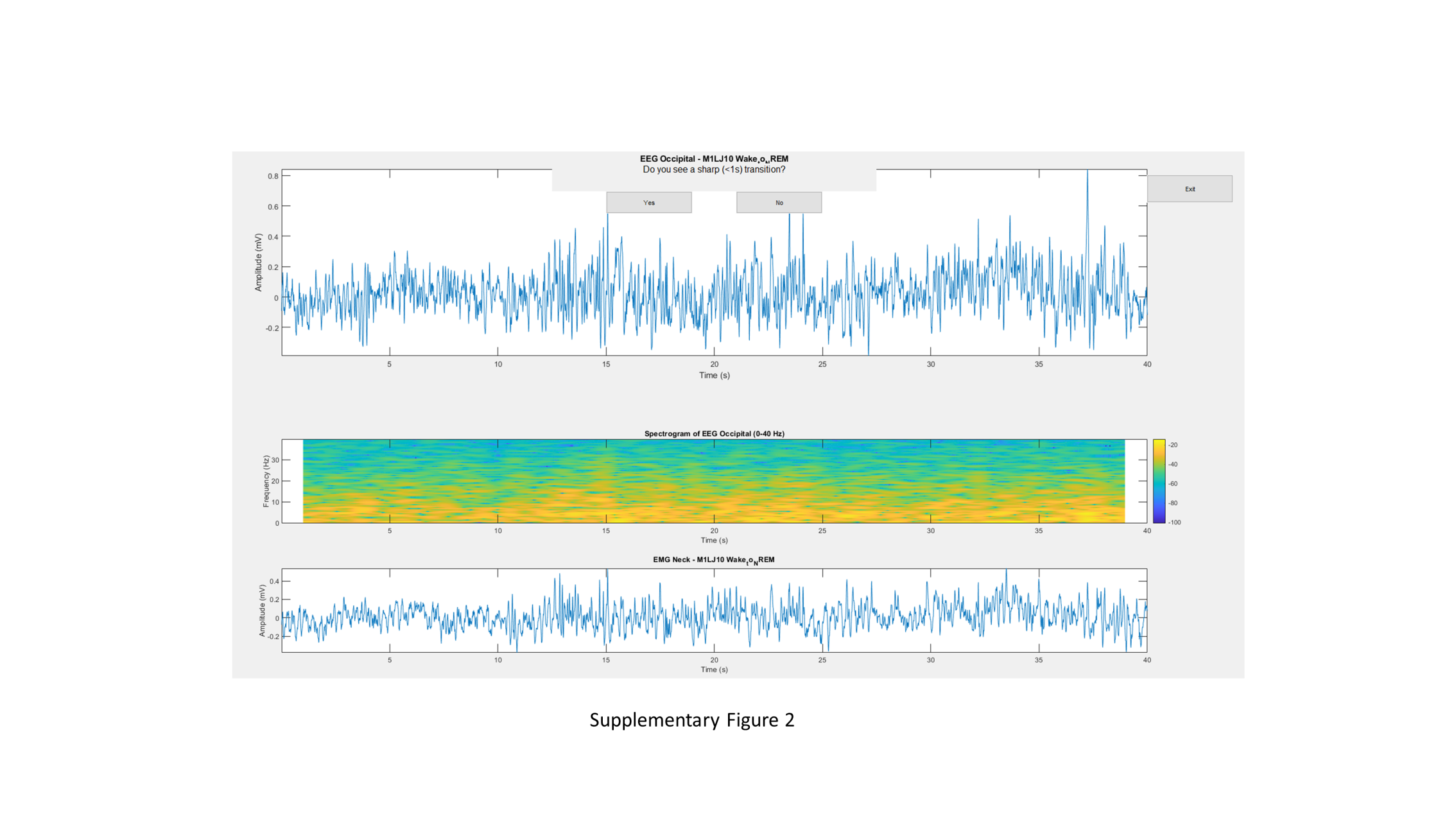
**Supplementary Figure 1. Custom built graphical user interface for annotation of vigilance state transitions.** (top panel) EEG recording from the left occipital cortex (middle panel) corresponding spectrogram and (bottom panel) simultaneously recorded EMG recorded in the nuchal muscle.

To avoid anchoring effects, each 56-s window was randomly shifted by up to ±16 s before presentation Experts and window of 40 seconds were presented to experts. The data shown is a representative sample intended to illustrate GUI functionality.

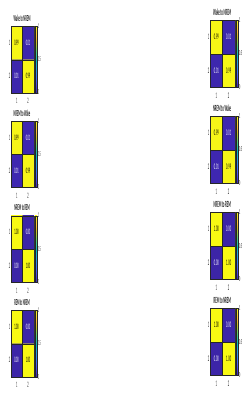

**Supplementary Figure 2. Confusion matrices for prediction of stable vigilance states in training dataset.**
